# Molecular mechanisms underlying the determination of axillary bud fate and outgrowth into branch crown in strawberry

**DOI:** 10.1101/2025.03.27.645709

**Authors:** Marie Alonso, Pierre Prévost, Aline Potier, Pascal GP Martin, Yves Caraglio, Michael Nicolas, Michel Hernould, Christophe Rothan, Béatrice Denoyes, Amèlia Gaston

**Affiliations:** Univ. Bordeaux, INRAE, Biologie du Fruit et Pathologie, UMR 1332, F-33140 Villenave d’Ornon, France; CIRAD, UMR AMAP and Université de Montpellier, F-34398 Montpellier, France; Plant Molecular Genetics Department, Centro Nacional de Biotecnología-CSIC, Campus Universidad Autónoma de Madrid, 28049 Madrid, Spain

**Keywords:** Axillary bud, *Branched 1 (BRC1)*, fate, *Fragaria vesca*, outgrowth, sexual and asexual reproduction, transcriptome

## Abstract

▪ In strawberry, the axillary bud (AXB) can produce either an elongated stem called stolon giving a daughter-plant (asexual reproduction) or an inflorescence-bearing branch crown (BC) (sexual reproduction). The fate of the AXB depends on node position on the axis and on genetic and environmental factors. Here, in *Fragaria vesca*, we addressed the largely unanswered question of how molecular factors determine AXB fate.
▪ To get insights into the mechanisms already at play in a morphologically indistinguishable (undifferentiated) AXB, depending on its fate, we combined (1) the phenotypic characterization of AXB development throughout plant growth with (2) the RNA-seq analysis of undifferentiated AXBs, using three different genotypes producing either BCs or stolons (*fvetfl1* and *fvega20ox* mutants, *FveFT3* overexpressor).
▪ Results allowed the identification of genes regulating AXB fate and outgrowth, among which *FveBRC1*. The analysis of *FveBRC1* expression in genotypes combining various traits (perpetual/seasonal flowering; runnering/runneless) and the generation of CRISPR/Cas9 *brc1* mutants further demonstrated that *FveBRC1* plays a central role in the determination of AXB fate in strawberry, in addition to its well-known function in BC outgrowth.
▪ These original results provide new insights into the determination of AXB fate and, consequently, the control of fruit yield in strawberry.

## Introduction

Plant architecture, defined as the spatial organization of the plant, has been a major domestication trait and a key target for crop improvement target due to its significant impact on yield (Eshed and Lippman, 2019). It depends on the ontogenetic dynamics of both shoot apical meristem (SAM) and axillary meristems (AXM) (Barthélémy and Caraglio, 2007; Labadie *et al*., 2023a). During plant growth, SAM produces stem and initiates leaves with, in their axils, the axillary bud (AXB) containing AXM with leaf primordia. AXB can remain dormant or develop into a lateral branch along the primary shoot. In addition to lateral branches, AXB can also produce floral structures and specialized vegetative organs including those involved in asexual reproduction such as stolon (Wang *et al*., 2018; Guo *et al*., 2021). Stolons are diageotropic elongated stems, underground in potato where they produce tubers, aerial in strawberry where they produce daughter-plants (Zierer *et al*., 2021). In cultivated strawberry (*Fragaria x ananassa*), a major horticultural species, the balance between stolon and branch formation is of major importance as it determines fruit and daughter-plant yields (Tenreira *et al*., 2017). Strawberry exhibits a sympodial branching: after floral initiation under short-day conditions the previous year, the SAM transitions to a floral meristem in spring, and the uppermost AXB of the stem (primary crown) develops into inflorescence-bearing branch (extension crown) (Labadie *et al*., 2023a). AXBs can follow two possible fates: they can form either stolons or lateral branches (branch crowns, BC) (Savini *et al*., 2008; Tenreira *et al*., 2017; Labadie *et al*., 2023a,b). Once the AXB fate is determined, the stolon develops rapidly, whereas BC development begins as a dormant bud (non-extended branch crown) with dormancy period of variable length before outgrowth (Labadie *et al*., 2023a,b; Han *et al*., 2024). Unraveling the regulatory network governing AXB fate determination and outgrowth is therefore crucial for modulating the balance between sexual and asexual reproduction, ultimately impacting both fruit and plant yields (Gaston *et al*., 2021; Andrés and Koskela, 2022).

Despite its importance, how the AXB fate is determined, prior to subsequent differentiation into a stolon, or dormant bud and BC, has been little studied to date. In strawberry, AXB fate is governed by genetic factors and environmental conditions (Andrés *et al*., 2021; Andrés and Koskela, 2022), node position on the axis (Labadie *et al*., 2023a,b) and hormones including gibberellins (Hytönen *et al*., 2009; Tenreira *et al*., 2017). In addition to these factors, several genetic alterations in *F. vesca* can be harnessed to better understand the determinism of AXB fate in strawberry. We showed recently that the *FLOWERING LOCUS T3* (*FveFT3)* gene promotes branching when overexpressed in *F. vesca*, likely by altering AXB fate: instead of producing a stolon, the AXB develops directly into a BC, bypassing the dormancy phase before outgrowth (Gaston *et al*., 2021). *FveFT3* is a member of the CENTRORADIALIS/TERMINAL FLOWER 1/SELF-PRUNING (CETS) family (FT/TFL1) primarily known for its crucial role in flowering, with FT as a florigen and TFL1 as a floral repressor (Wickland and Hanzawa, 2015; Gaston *et al*., 2021). The interplay between FT and TFL1 is essential in shaping architecture (Moraes *et al*., 2019; Colleoni *et al*., 2024), including in strawberry (Gaston *et al*., 2021; Andrés and Koskela, 2022). In *F. vesca*, a natural mutation in *FveTFL1* leads to perpetual flowering, which is the ability to flower continuously along vegetative development through the sustained production of new BCs (Iwata *et al*., 2012; Koskela *et al*., 2012). Two other mutations influence AXB fate, a mutation in the *FveGA20ox4* gibberellin (GA) biosynthesis gene responsible for the natural runnerless (no stolon) phenotype (Tenreira *et al*., 2017; Andrés *et al*., 2021), whereas a mutation in the GA signaling gene, *FveRGA1* (*REPRESSOR OF GIBBERELLIC ACID1*) of the *DELLA* family, reversing the effect of the *fvega20ox4* mutation (Caruana *et al*., 2018; Li *et al*., 2018). Despite these findings, the regulatory network and molecular mechanisms controlling AXB fate determination in strawberry remains largely unexplored

The regulation of AXB outgrowth into lateral branches in plants is comparatively better documented. AXB outgrowth has been well-studied in species producing only lateral branches, where AXB activation from dormancy to lateral branch outgrowth is mainly controlled by the apical dominance exerted by SAM (Cline, 1997; Barbier *et al*., 2019) and crosstalk between the plant hormones auxin, cytokinin, strigolactone and abscisic acid (Ferguson and Beveridge, 2009; Wang *et al*., 2018; Barbier *et al*., 2019). Stolon-bearing species are more complex to study as AXB outgrowth results in either stolon or BC. In octoploid strawberry, the GATA transcription factor *HANABA TARANU (FaHAN)* has been identified as a regulator of AXM initiation and AXB outgrowth into stolon (Liang *et al*., 2022) while in the diploid strawberry *FveMYB117a* inhibits AXB outgrowth into BC by regulating the cytokinin pathway (Han *et al*., 2024). The regulatory network controlling AXB outgrowth into a lateral branch converges on a major repressor which interacts with FT/TFL1, the *TCP (TEOSINTHE BRANCHED 1/CYCLOIDEA/ PROLIFERATING CELL FACTOR)* transcription factor, *BRANCHED1 (BRC1)* (Aguilar-Martinez *et al*., 2007; Wang *et al*., 2019; Colleoni *et al*., 2024). Based on the known role of *BRC1* in AXB dormancy and outgrowth into lateral branches, and its expression pattern in strawberry, the closest *F. vesca* homolog of *AtBRC1*, *FveTCP9* (hereafter referred to as *FveBRC1*), was hypothesized to play a role in AXB outgrowth into BC (Wei *et al*., 2016; Qiu *et al*., 2019; Han *et al*., 2024).

Here, we reasoned that analyzing undifferentiated AXBs while considering their subsequent fate would provide a comprehensive understanding of the molecular mechanisms of AXB fate determination. To this end, we first characterized early AXB development in *F. vesca* to establish a developmental scale describing the stages of AXB outgrowth. We then used this scale to examine the spatio-temporal growth of AXBs along the primary crown according to their subsequent fates, stolon or BC. Next, we performed a transcriptome analysis of morphologically undifferentiated AXBs with distinct predicted fates to uncover early molecular signatures underlying AXB determination. Finally, we investigated *FveBRC1*, highlighted through transcriptome analysis, revealing its pivotal and original role in AXB fate determination, in addition to its already known role in AXB activation and BC outgrowth. These findings make *FveBRC1* a promising target for improving cultivated strawberries.

## Materials and methods

### Plant material and growth conditions

Diploid strawberry (*Fragaria vesca*) genotypes were used in this study. Seeds were sown in a mixture of two-thirds loam and one-third grit and grown in long day conditions (16-h/8-h day/night at 22°C/18°C; 250 μmol m^−2^ sec^−1^). Plants were transplanted into 1-liter pots containing the same mixture and were maintained in a greenhouse under natural conditions. Three genotypes carrying the *tfl1* mutation that produces the perpetual flowering [PF] phenotype (Iwata *et al*., 2012) were used for studying AXB development and plant architecture as well as for transcriptome analysis. These genotypes included: ‘Hawaii-4’ (H4); the *35S::FveFT3FveOE* (*FT3^OE^*) transgenic line, which overexpresses *FveFT3* in the H4 genetic background (Gaston *et al*., 2021); and the runnerless [r]/[PF] ‘Reine des Vallées’ (RdV) variety, which is a *FveGA20ox4/FveTFL1* mutant (Tenreira *et al*., 2017). H4 was used for gene editing with the CRISPR/Cas9 system (CR-*fvebrc1* mutants). To analyze *FveBRC1* expression in different mutant backgrounds, we generated four *F. vesca* genotypes combining the seasonal flowering [SF]/perpetual flowering [PF] and the runnering [R]/runnerless [r] phenotypes in a near-isogenic background. These genotypes, derived from an ‘Ilaria’ F1 hybrid heterozygous for both *FveGA20ox4* and *FveTFL1* previously obtained by crossing the ‘Alpine’ *ga20ox4/tfl1* mutant with ‘Sicile’ (Tenreira *et al*., 2017), were brought to homozygosity by five successive selfings (S5) and therefore named ‘Ilaria’ S5-1 to S5-4.

### Plant dissection for the determination of AXB development and plant architecture

Strawberry AXBs were observed by dissecting two- to four-month-old seedlings using a stereomicroscope (SZX16; Olympus, Tokyo, Japan). The AXB developmental scale was established by analyzing the spatio-temporal development of AXBs using photographs obtained with the EPview software driving an Olympus EP50 camera mounted on the stereomicroscope. Each description of an AXB included its developmental stage and node position (Labadie *et al*., 2023a). The plant developmental stage was determined according to the number of developed trifoliate leaves and of newly non-expanded leaves (Fig. S1). Patterns of AXB development were determined on 10 plants per genotype, starting from the 3-leaf stage up to flowering. In Ilaria-S5 lines, 5 to 8 plants of each of the four genotypes were dissected after three months of growth under long-day conditions following sowing. For CR-*fvebrc1* mutants, the number of trifoliate leaves, crowns, stolons and inflorescences were counted on 2- to 18-months-old primary transformants. Architecture of all genotypes was described after dissecting plants as described in Gaston *et al*. (2021).

### Histological studies and in situ hybridization

For histological studies and in situ hybridization, longitudinal sections of the primary crown were prepared from 3-5 trifoliate leaf seedlings after leaf removal. Dissected samples were prepared as described in Tenreira *et al*. (2017). For in situ hybridization of *FveBRC1*, undifferentiated AXBs were collected from 3 months-old plants of the H4, *FT3^OE^* and RdV genotypes. ISH-*FveBRC1* primers used for synthesizing digoxygenin-UTP-labelled sense and antisense RNA probes are provided in Table S1.

### RNA-seq and qPCR analyses

For RNA-seq and qPCR analysis, undifferentiated AXBs were collected at the fourth node of 4-leaf stage plants on the newly non-expanded leaf on studied genotypes, immediately frozen in liquid nitrogen and stored at -80°C until analysis. Each genotype was represented by three biological replicates. Each sample was constituted by 30 AXBs comprising the AXM and one or two leaf primordia collected on 30 plants. Total RNA was extracted as described in Gaston *et al*. (2021). For RNAseq analysis on H4, *FT3^OE^* and RdV genotypes, stranded mRNA-seq libraries were prepared and sequenced (150 bp paired-end reads) by GenoToul (Toulouse, France). The *F. vesca* v4.0.a1 genome (Edger *et al*., 2017) and its v4.a2 annotation (Li *et al*., 2019a) were used as reference. Reads were mapped using STAR v2.7.9a (Dobin *et al*., 2013) with default parameters. Gene level draw counts and Transcripts per Million (TPM) values were obtained with RSEM (v1.3.3) (Li and Dewey, 2011). Differential expression was analyzed using the ‘exactTest()’ function of the R/Bioconductor edgeR package (v3.32.1) following Relative Log Expression (RLE) normalization and multiple testing correction with the Benjamini–Hochberg procedure. Gene expression changes were considered significant using a cutoff of false discovery rate (FDR) 0.05 and an absolute log2 (fold-change) greater than 1. Exploratory analysis was conducted using the R software (v3.6.1). Principal component analysis (PCA) was performed on TPM values using the 500 most variable genes with the ‘FactoMineR (v2.4)’ and ‘factoextra (v1.0.7)’ packages. Venn diagrams were created using the Venny 2.1.0 platform (BioinfoGP, CNB-CSIC). Hierarchical clustering of genes differentially expressed in at least one comparison and heatmaps were obtained from z-scores calculated from TPM values using ‘EnrichedHeatmap (v1.16.0)’ package. Fisher’s exact tests (R package ‘TopGO’ v2.38.1) were used to identify the Gene Ontology Biological Processes enriched in each cluster. Gene Set Enrichment Analysis (GSEA) was carried out with the ‘ClusterProfiler’ package (v4.12.6). The ‘Dormancy’, ‘Bud activation’, ‘BRC1up’ and ‘BRC1down’ genes sets were previously described (Nicolas *et al*., 2022; Tarancón *et al*., 2017). Parameters for studying the overrepresentation of the different gene sets were False Discovery Rate (FDR) for method and 0.5 for pvaluecutoff. Quantitative RT-PCR was performed following the protocol described in Gaston *et al*. (2021). Primers are provided in (Table S1).

### Plasmid constructs and plant transformation

CRISPR/Cas9 mutagenesis of *FveBRC1* using the H4 genotype was carried out with two sgRNAs (Table S1) as described in Gaston *et al*. (2021). Resulting mutations were detected by sequencing T1 lines. Plants carrying four independent homozygous mutations and a heterozygous mutation were selected for further analysis.

### Statistical analyses

Pairwise multiple comparisons were carried out on Ilaria population and *CR-fvebrc1* gene expression results with Kruskal-Wallis test (ANOVA) followed by a Tukey post hoc test. Wilcoxon-Mann-Whitney test was used for phenotypic analysis to compare the number of leaves, stolons, BCs and inflorescences in WT and *CR-fvebrc1* mutants.

## Results

### Establishment of an AXB developmental scale

To provide a detailed description of the early stages of AXB development, three genotypes producing either stolons (‘Hawaii-4’ (H4)) or BCs (*FT3^OE^*and ‘Reine des Vallées’ (RdV)) (Fig. 1a,b) were characterized at morphological (Fig. 1c-i) and histological (Fig. 1j-p) levels. In all three genotypes, the AXB in the axil of the newly non-expanded leaf was triangular in shape, with two prophylls protecting the AXM, which displayed one or two primordia (H4 and *FT3^OE^*; Fig. 1c,j,f,m and RdV; Fig. S2). Elongation of the first internode occurred first at the stoloniferous bud apex (Fig. 1k). Later in stolon development, we observed the elongation of the second internode and the development of leaves in the daughter plant (Fig. 1l). In *FT3^OE^* and RdV, the differentiation into BC was associated with the thickening of the AXB (Fig. 1g-n; Fig. S2). Then, the first leaf grew beyond the top of the prophylls (Fig. 1h-o) until the leaf was fully developed (Fig. 1i,p).

**Fig. 1.**
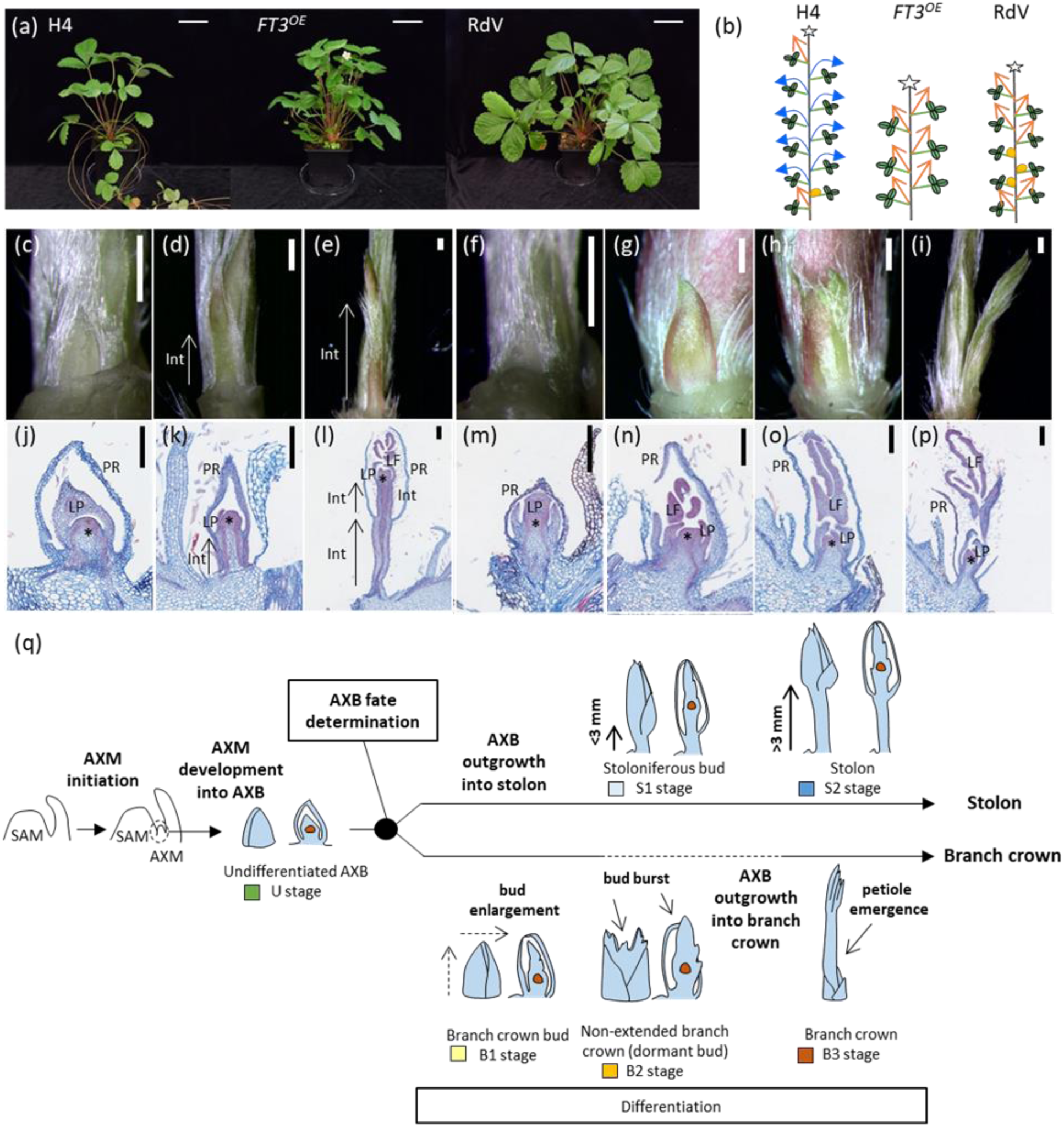
Developmental stages of the axillary bud (AXB) fate determination and differentiation into stolon or branch crown (BC). (a) ‘Hawaii-4’ (H4), *35S::FveFT3^FveOE^* (*FT3^OE^*) and ‘Reine des Vallées’ (RdV) 4-month-old plants. White bar, 5 cm. (b) Schematic plant architecture of the primary crown of each genotype. Each primary crown (green vertical line) is terminated by an inflorescence (white star). The AXB in the axil of each leaf can differentiate into a stolon (blue arrow), a branch crown (BC) (orange arrow), or remain dormant (orange circle). (c-i) Morphological characterization of AXB differentiation into a stolon or a BC. White bar, 500 µm. (c-e) Differentiation of AXB into a stolon in H4. The white arrow indicates the first internode (Int), which elongates in (d) stoloniferous bud and (e) stolon. (f-i) Differentiation of AXB into a BC in *FT3^OE^*. (j-p) Histological characterization of AXB differentiation into stolon or BC. AXB sections were stained with astra bleu and safranin for differentiated and dividing cells, respectively. The AXB, surrounded by prophylls (PR), shows a meristematic dome (black asterisk) at its center and developing leaf primordia (LP). LP develop into leaves (LF, leaf). In AXB differentiating into stolon (j-l), the black arrow in (k) indicates the first internode (Int), the elongation of the stoloniferous bud and (l) the elongation of the two first internodes of the stolon. (m-p) AXB differentiation into a BC. Black bar, 200µm. (q) AXB development scale starting with axillary meristem (AXM) initiation, followed by AXM development into undifferentiated AXB (U stage), and AXB decision to differentiate into either a stolon or a BC (AXB fate determination). The AXB differentiation into stolon is characterized by the elongation of the first internode, defined as the stoloniferous bud stage (S1, elongation <3mm), which is followed by the stolon stage (S2, elongation >3mm). The differentiation of AXB into BC is characterized by the enlargement of the AXB (B1 stage), followed by a dormancy period (dormant bud or non-extended BC, B2 stage), and by AXB outgrowth into BC that is characterized by petiole emergence (B3 stage). Each stage of AXB development is characterized by a different color as described in the legend of Figure 2.

These observations were used to establish a scale for AXB development (Fig. 1q). The first stage was observed in the axil of the newly non-expanded leaf. At this stage, AXBs were morphologically indistinguishable regardless of their final fate, stolon or BC. We have therefore called this stage the undifferentiated stage (U stage), during which the two different developmental fates (stolon or BC) are likely already determined. For the AXB differentiating into stolon, we set a threshold for first internode elongation at 3 mm, below which the AXB was considered as a stoloniferous bud (S1 stage) and above which it was considered as a stolon (S2 stage). When the AXB produced a BC, the thickened bud was named BC bud (B1 stage). When the bud burst and the leaf began to be visible, we considered AXB to be a non-extended BC (dormant bud, stage B2); the emergence of a leaf petiole from the BC bud indicated the BC development (B3 stage).

### AXB fate depends on its node position on the primary crown and the genotype

To characterize the spatio-temporal dynamics of AXB outgrowth into a stolon or BC, we determined the plant architecture of the three genotypes, H4, *FT3^OE^* and RdV, starting from the 3-leaf stage (i.e. two developed leaves plus a new, non-expanded leaf) to the 6-leaf and floral stages (Fig. 2). We then scored AXB stages at each node on the primary crown, according to the established scale (Fig. 1q). For the three genotypes, undifferentiated AXBs were observed only in the axil of non-expanded leaves, which corresponded to the last-emerged node. Below this point, AXBs were already differentiated, displaying a gradient of developmental stages, with those closer to the SAM exhibiting a less advanced stage.

**Fig. 2.**
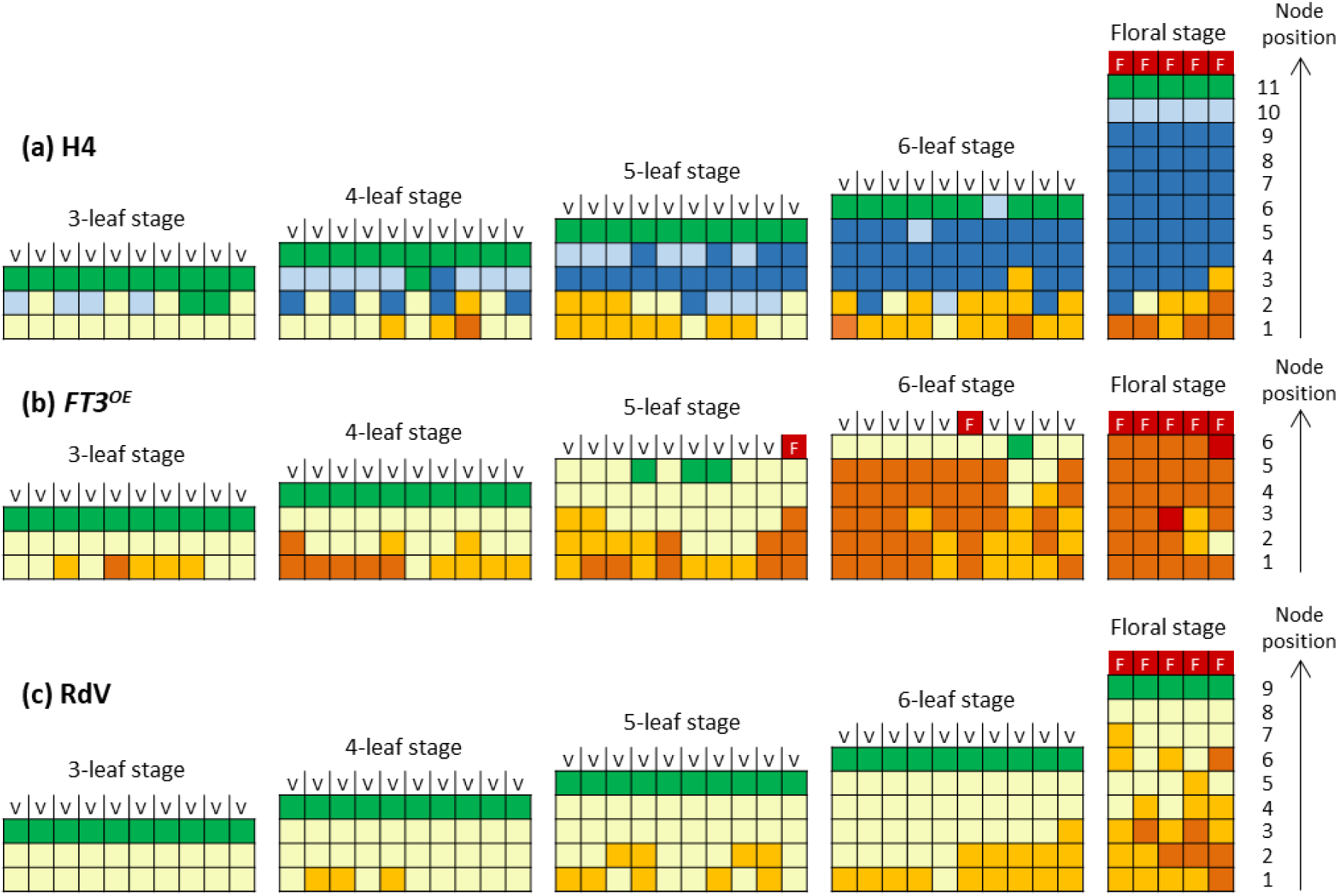
Patterns of AXB development from the vegetative stage to flowering in (a) ‘Hawaii-4’ (H4), (b) *35S::FveFT3^FveOE^* (*FT3^OE^*) and (c) ‘Reine des Vallées’ (RdV) plants. The architecture of ‘Hawaii-4’ (H4), *35S::FveFT3^FveOE^* (*FT3^OE^*) and ‘Reine des Vallées’ (RdV) plants was determined by dissection of plants at the 3- to 6-leaf stages (n = 10) to the flowering stage (n = 5). At a given stage (X) there are X-1 developed leaves and one emerging leaf. Each column represents a single plant. Each square box represents a node including the AXB. Box colors correspond to the stages of AXB development as described in Figure 1 legend: green, undifferentiated bud (U); pale blue, stoloniferous bud (S1); blue, stolon (S2); yellow, BC bud (B1); orange, dormant bud or non-extended branch crown (B2); dark orange, branch crown BC (B3). Red box, floral BC. V = vegetative apical meristem, F = floral apical meristem.

The AXB fate was dependent on the genotype. In the H4 genotype, AXBs at nodes 3 to 9 developed into stolons (S2 stage) (Fig. 1b; Fig. 2a) while AXBs of the first two nodes could eventually produce BCs. In the *FT3^OE^* and RdV genotypes, all AXBs differentiated into BC buds (B1 stage) (Fig. 1b; Fig. 2b,c). AXB development into BCs was rapid in *FT3^OE^*, with the production of vegetative branches (B3) starting at the 3-leaf stage. In RdV, AXB outgrowth progressed more slowly, reaching the B2 stage; some B2 AXBs, particularly those located at nodes 1 to 3, could develop into BCs (B3 stage) after a variable period of dormancy (Fig. 2b,c).

Flowering time was also variable between the genotypes. In RdV and H4, flowering took place at the 9- and 11-leaf stages respectively, while *FT3^OE^* flowered earlier, at the 6-leaf stage (Fig. 2). In RdV and H4, the uppermost AXB of the primary crown produced an inflorescence-bearing branch (extension crown) after flowering (Fig. 2a,c), thus enabling sympodial branching (Fig. 1b). Remarkably, thanks to the detailed characterization of the temporality of plant development for the three genotypes (Fig. 2), we were able to predict with great confidence the AXB fate while they were still undifferentiated. From the fourth to the penultimate AXB, we could determine that the final AXB fate was the production of a stolon in H4 and BC in RdV and *FT3^OE^*, with or without an intermediate step of dormancy, respectively.

### Transcriptome analysis of undifferentiated AXBs provides comprehensive pictures of the determination of AXB fate and regulation of branch crown formation by *FveFT3*

To evidence the early molecular programs implemented in the undifferentiated AXB, we collected undifferentiated AXBs with known fates (stolon or BC, with or without AXB dormancy) at node 4 on 4 leaf-stage plants of the three genotypes studied (Fig. 3a) and subjected them to transcriptome analyses. Principal component analysis (PCA) enabled us to clearly discriminate between the three genotypes (Fig. 3b). The first two dimensions (PC1 and PC2) explained 54% and 16% of the structural variance, respectively. PC1 clearly separated RdV from H4 and *FT3^OE^*, thus highlighting the strong effect of the genetic background. PC2 allowed the distinction of all three genotypes with a clear separation between H4 and *FT3^OE^*. Both share the same genetic background, indicating the strong effect of *FveFT3* overexpression on gene expression as early as the undifferentiated AXB stage.

**Fig. 3.**
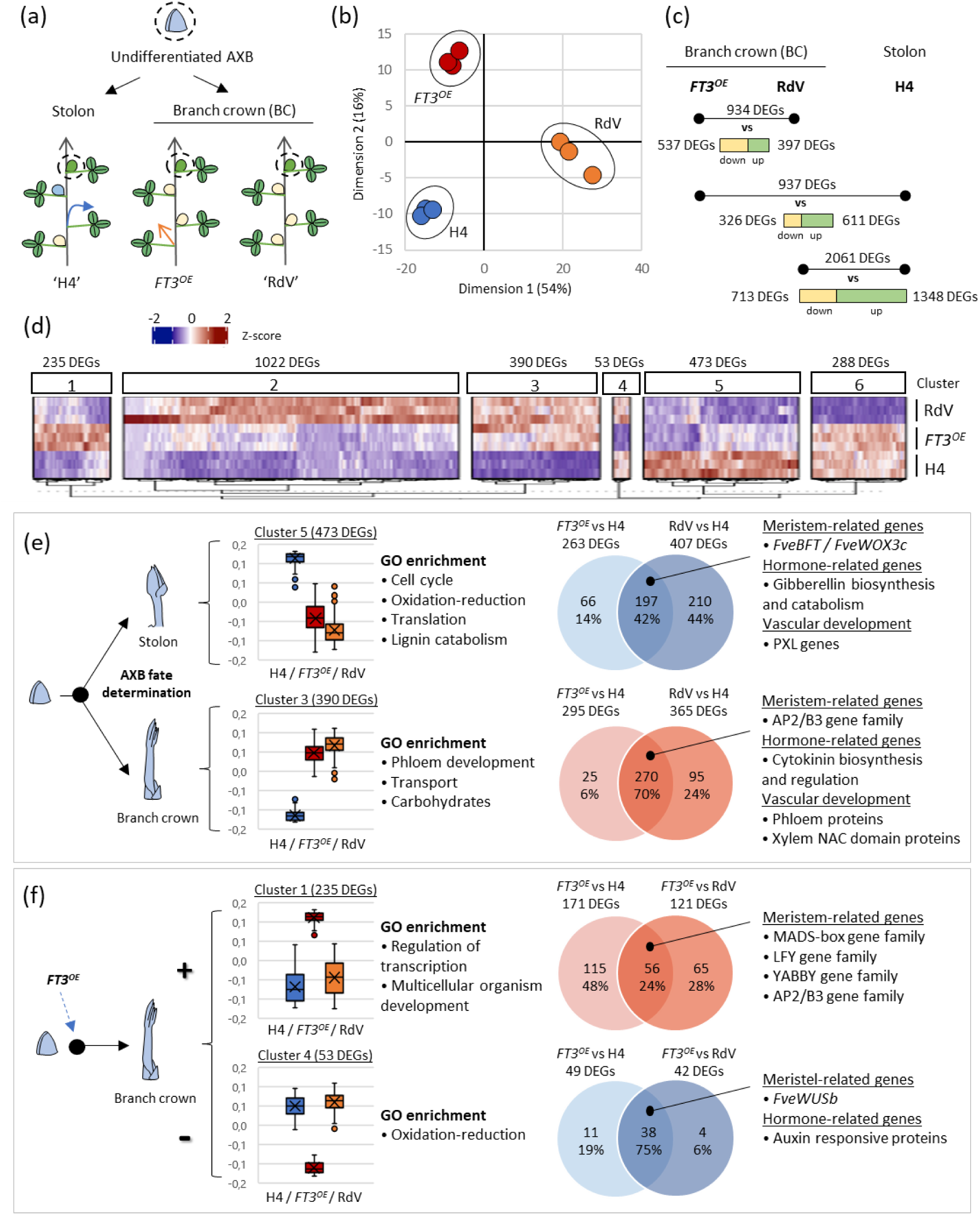
Differential gene expression analysis between undifferentiated AXBs of ‘Hawaii-4’ (H4), *35S::FveFT3^FveOE^*(*FT3^OE^*) and ‘Reine des Vallées’ (RdV) genotypes. (a) RNAseq design. Dotted circles represent the undifferentiated AXBs collected from node 4 of plants at the 4-leaf stage of the three genotypes H4, *FT3^OE^*and RdV. AXBs of H4 differentiate into a stolon while AXBs of *FT3^OE^*and RdV differentiate into a branch crown (BC). (b) Principal component analysis (PCA) plot of the 500 genes with the most variable expression. (c) Total number of significant differentially expressed genes (DEGs) in each pairwise comparison (*FT3^OE^ vs* RdV, *FT3^OE^ vs* H4 and RdV *vs* H4). Green and yellow bars indicate the number of up-regulated and down-regulated genes, respectively, for each pairwise comparison. (d) Hierarchical clustering analysis (HCA) of the 2461 DEGs identified in at least one pairwise comparison between the 3 genotypes. Each column represents the normalized expression values (z-score) of a transcript for each sample in a row (including triplicates). Red indicates a high transcript abundance, blue low. Six clusters were identified according to the degree of similarity of the expression profile of each gene for the three genotypes. (e) DEGs from cluster 3 and 5 involved in the determination of AXB fate and (f) DEGs from cluster 1 and 4 involved in the regulation by *FveFT3* of AXB differentiation into BC and BC outgrowth. Boxplots show gene expression patterns (triplicates average values of z-scores) for the three genotypes. Selected representative GO terms enriched in each cluster are listed. Venn diagrams represent the number of DEGs and common DEGs in the pairwise comparisons indicated. Selected categories of genes enriched in common DEGs are listed.

To investigate at the molecular level the determination of AXB fate, regulation of BC differentiation and development by *FveFT3* and induction of dormancy, we next performed pairwise comparisons of differentially expressed genes (DEGs) between AXBs of the three genotypes. Comparison of AXB transcriptomes revealed DEGs between genotypes with (i) a different AXB fate and the same genetic background (*FT3^OE^ vs* H4; 937 DEGs), (ii) a same AXB fate but a different genetic background (*FT3^OE^ vs* RdV; 934 DEGs) and (iii) a different AXB fate and genetic background (RdV *vs* H4; 2061 DEGs) (Table S2) (Fig. 1b; Fig. 3c). The highest change in total number of DEGs was observed between H4 and RdV, which is consistent with their different genetic background and AXB fate (Fig. 3c).

To identify groups of genes sharing similar expression patterns, we performed hierarchical clustering) of the 2461 DEGs identified in at least one pairwise comparison and identified six clusters (C1 to C6) (Fig. 3d; Table S2). In clusters 3 and 5 (390 and 473 DEGs, respectively), the transcript profile of H4 (stolon) is opposite to those of *FT3^OE^* and RdV (BC) suggesting clusters involved in AXB fate determination. In cluster 1 and 4 (235 and 53 DEGs, respectively), the transcript profile of *FT3^OE^* is opposite to those of RdV and H4 suggesting a specific regulation of AXB differentiation by *FveFT3*. In clusters 2 and 6 (1023 and 288 DEGs, respectively), the transcript profile of RdV (dormant AXBs) is opposite to those of H4 and *FT3^OE^* (non-dormant AXBs), suggesting gene expression patterns associated with AXB dormancy; although, an effect of the genetic background (RdV vs H4/ *FT3^OE^*) cannot be ruled out.

### Identification of key genes involved in the AXB fate determination

To identify genes positively associated with AXB determination into stolon, we examined the 473 DEGs in cluster 5, which exhibited higher transcript levels in H4 (stolon) than in *FT3^OE^* and RdV (BC). Gene Ontology (GO) enrichment analysis indicated that it was enriched in morphogenesis-related terms likely reflecting early cell growth processes such as ‘protein translation’ and ‘cell cycle’ as well as later specialization such as ‘photosynthesis’, ‘anatomical structure development’ and ‘lignin catabolism’ (Fig. 3e; Table S3). Interestingly, we found three *TCP* genes: *FveBRC1*, the closest homologue to *AtBRC1*, a major repressor of shoot branching by promoting AXB dormancy (Aguilar-Martinez *et al*., 2007; Wang *et al*., 2019), and *FveTCP18* and *FveTCP11*, belonging to the PCF clade (Fig. S3). Among the 473 cluster 5 DEGs, we then focused on the 197 DEGs common to the *FT3^OE^*vs H4 and RdV vs H4 comparisons (Fig. 3e). These 197 DEGs strictly associated with AXB determination into stolon were involved in ‘transcriptional regulation’, ‘hormone biosynthesis and signaling’, and ‘vascular development’ (Table S3). These 197 DEGs included the meristem-related genes *BROTHER OF FT AND TFL1* (*FveBFT)* and *WUSCHEL-related HOMEOBOX3c (FveWOX3c)*; a set of genes involved in hormone biosynthesis and signaling (GA, ABA, auxin, ethylene), including one GA biosynthesis gene (*FveGA20ox4*), four GA metabolism genes (*FveGA20ox4, FveGA2ox4, FveGA20ox1,* and *FveGA2ox1*), as well as the ABA receptor-encoding gene *PYRABACTIN RESISTANCE-Like4 (FvePYL4)*; and the leucine-rich repeat receptor-like protein kinases, *PHLOEM INTERCALATED WITH XYLEM-LIKE (FvePXL1* and *FvePXL2)*, which regulate vascular-tissue development in the stem (Fig. 3e; Fig. 4a; Table S2).

**Fig. 4.**
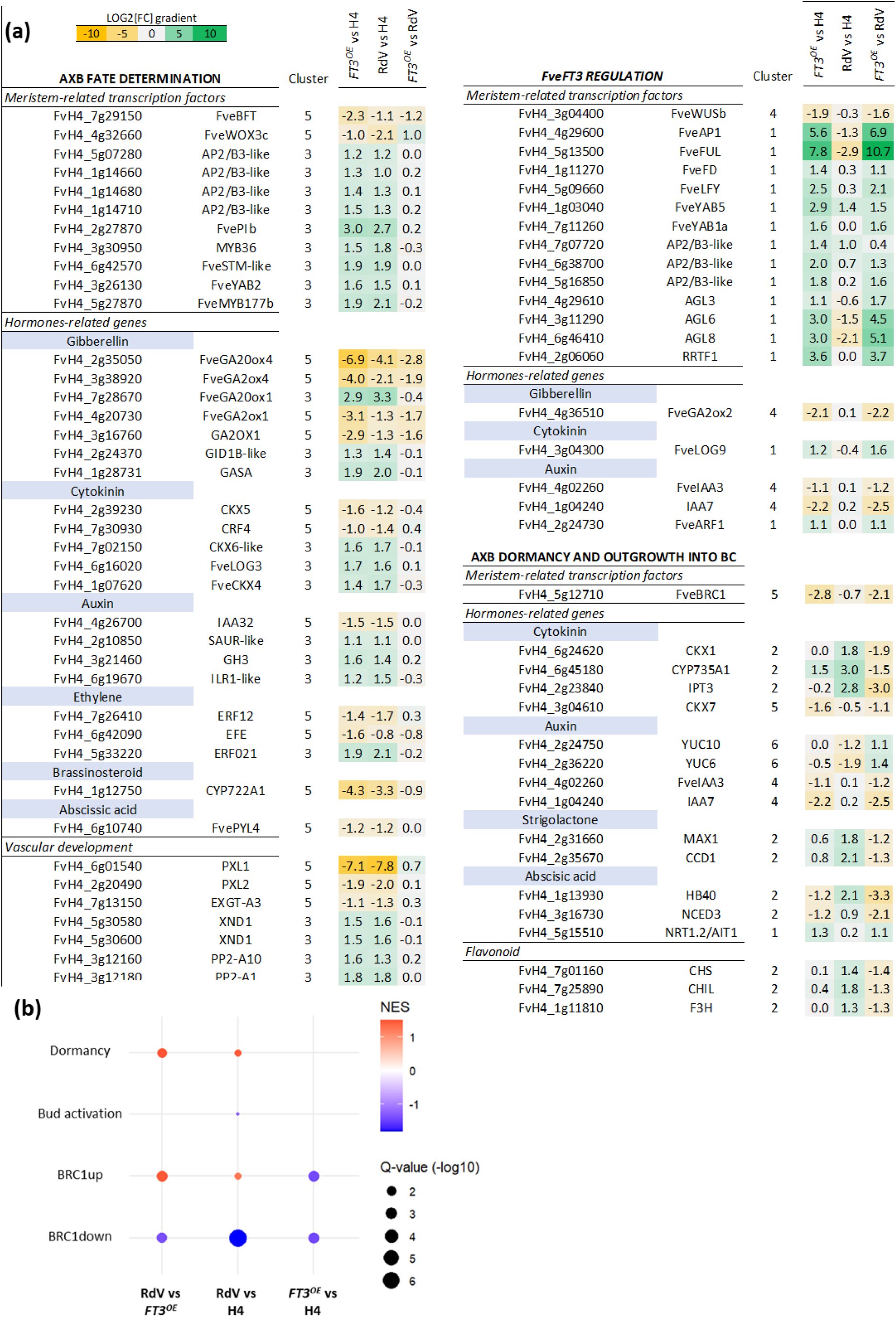
AXB fate, dormancy and outgrowth into branch crown (BC). (a) Heatmaps of DEGs associated with AXB fate, regulation by *FveFT3* and AXB dormancy and outgrowth into BC as determined by RNAseq analysis of undifferentiated AXBs of ‘Hawaii-4’ (H4), *35S::FveFT3^FveOE^*(*FT3^OE^*) and ‘Reine des Vallées’ (RdV) genotypes. Selected categories of genes are shown. (b) Gene Set Enrichment Analysis (GSEA) of genes issued from the RdV vs *FT3^OE^*, RdV vs H4, *FT3^OE^* vs H4 was performed using dormancy, bud activation, BRC1 dependent upregulated genes (BRC1up) and BRC1-dependent downregulated genes (BRC1down) gene sets.

To identify genes positively associated with AXB determination into BC, we next focused on the 390 DEGs of cluster 3 whose transcript abundances were higher in *FT3^OE^* and RdV (BC) than in H4 (stolon). GO enrichment analysis indicated that it was enriched in ‘phloem development’, ‘transport’ and ‘carbohydrates’ terms, thus highlighting the importance of the vascularization process and the requirement for energy in the BC formation (Fig. 3e; Table S3). Among cluster 3, 270 DEGs were common between *FT3^OE^* vs H4 and RdV vs H4 comparisons (Fig. 3e; Table S2). Common DEGs included transcription factors potentially involved in the regulation of meristem activity like *PISTILLATAb (FvePIb)*, *FveMYB117b*, *YABBY2 (FveYAB2), SHOOT MERISTEM LESS (STM)* and four *AP2/B3-like* genes; genes related to hormone (auxin, cytokinin, gibberellin, ethylene) biosynthesis and signaling, like the cytokinin pathway genes *CYTOKININ OXIDASE/DEHYDROGENASE* (*FveCKX4)* and *LONELY GUY3 (FveLOG3)* and the gibberellin pathway genes *FveGA20ox4* and *GA INSENSITIVE DWARF1 (FveGID1)*; and genes related to vascular development like two *PHLOEM PROTEIN* and two *XYLEM NAC DOMAIN* (Fig. 4a; Table S2). Altogether, these findings reflect the early specialization of AXBs which, depending on their fate as stolons or BCs, exhibit contrasted transcriptomes already at the undifferentiated stage.

### Identification of key genes involved in the regulation of branch crown differentiation by *FveFT3*

To further explore the role of *FveFT3*, we then analyzed clusters 1 and 4 (Fig. 3f). Overexpression of *FveFT3* in H4 provokes a bushy plant phenotype because *FveFT3* promotes AXB differentiation into a BC at the expense of stolon production, likely through the reprogramming of AXB (Gaston *et al*., 2021). Moreover, in *FT3^OE^*, the dormant bud (non-extended BC) stage is by-passed so that BCs develop rapidly from AXBs (Fig. 1q). GO enrichment analysis revealed that cluster 1 (overexpression in *FT3^OE^*) is enriched in regulation and morphogenesis-related terms including ‘regulation of transcription’ and ‘multicellular organism development’ and cluster 4 (downregulation in *FT3^OE^*) in the ‘oxidation-reduction’ term (Fig. 3f; Table S3). To select genes strictly linked to *FveFT3* overexpression, we further examined DEGs common to the *FT3^OE^* vs H4 and *FT3^OE^* vs RdV pairwise comparisons and found 56 and 38 common DEGs in cluster 1 and 4, respectively (Fig. 3f; Table S2). Interestingly, a large number of meristem-related transcription factors were up-regulated in *FT3^OE^* (cluster1) including flowering-related genes such as *LEAFY (FveLFY), FLOWERING LOCUS D (FveFD)* and fives MADS-box family genes like *FRUITFULL (FveFUL)*, *APETALA1 (FveAP1)*, and *AGAMOUS-like* (*AGL8*, *AGL6* and *AGL3*). Most of these MADS-box genes showed very high transcript abundance in *FT3^OE^*, the log2FC being, for example, higher than 10 for *FveFUL* in the *FT3^OE^* vs RdV comparison. Other meristem-related transcription factors were upregulated in *FT3^OE^* (cluster 1) such as two *YABBY* genes (*FveYAB5* and *FveYAB1a)* whereas *WUSCHELb (FveWUSb)* was down-regulated (cluster 4) (Fig. 3f; Table S2). Up-regulated hormone-related genes included ethylene-responsive transcription factor *Redox-Responsive Transcription Factor 1* (*RRTF1*), *Auxin Response Factor 1* (*FveARF1*) and cytokinin-related gene *LONELY GUY 9 (FveLOG9)* while down-regulated hormone-related genes included two *INDOLE-3-ACETIC ACID* genes *(IAA2 and IAA4*) and the gibberellin catabolism gene *FveGA2ox2* (*GA2 oxidase2*). These findings indicate that *FveFT3* overexpression not only triggers the expression of genes involved in the meristem activity necessary for transition to BC but also in the flowering, which occurs much later.

### Identification of key genes involved in the induction of AXB dormancy

To identify genes possibly involved in the induction of AXB dormancy, we focused on genes differentially expressed between *FT3^OE^* and RdV, both genotypes exhibiting a BC fate, without or with a dormancy period, respectively. DEGs negatively regulated in *FT3^OE^ vs* RdV are likely to be associated with AXB dormancy induction, while those positively regulated could be involved in AXB activation and outgrowth into BC (B2 to B3 stage; Fig. 1q; Fig. 4a). We chose to focus on genes with known roles in shoot branching/dormancy induction (Rameau *et al*., 2015, Wang *et al*., 2018, Barbier *et al*., 2019). Not surprisingly, *FveBRC1*, known as a major repressor of shoot branching and promoting AXB dormancy in other species (Wang *et al*., 2019) was down-regulated in *FT3^OE^*. Downregulated DEGS also included hormone-related genes involved in ABA biosynthesis and signaling (*NINE-CIS-EPOXYCAROTENOID DIOXYGENASE 3, NCED3; HOMEOBOX PROTEIN 40, HB40*), cytokinin biosynthesis (*ISOPENTENYL TRANSFERASE, IPT3;* CYTOKININ OXIDASE/DEHYDROGENASE, *CKX; CYTOCHROME P450*, *CYP735A1*), auxin signaling (*INDOLE-3-ACETIC ACID, IAA3 and IAA7)*, and strigolactone biosynthesis (*CAROTENOID CLEAVAGE DIOXYGENASE, CCD1; MORE AXILLARY BRANCH, MAX1*). They also included several flavonoid biosynthetic genes (*CHALCONE SYNTHASE, CHS; CHALCONE ISOMERASE, CHI; FLAVANONE 3-HYDROXYLASE, F3H*) that are potentially involved in the regulation of AXB outgrowth under the control of *MAX1* (Fig. 4a) (Lazar *et al*., 2006). Fewer genes including the auxin biosynthesis genes *YUCCA (YUC6 and YUC10)* and the *ABA-IMPORTING TRANSPORTER (NRT1.2/AIT1)* were upregulated in *FT3^OE^*. These findings, which indicate the downregulation in *FT3^OE^* of *FveBRC1* and its downstream targets, such as HB40 and NCED3 (Gonzalez-Grandio *et al*., 2017), highlight the prominent role likely played by *FveBRC1* in regulating strawberry AXB dormancy and activation. Remarkably, *FveBRC1*, which belongs to cluster 5, is less expressed in *FT3^OE^* and RdV than in H4 (Fig. 4a), suggesting a dual role for this gene, first in the determination of AXB fate and, later, in the regulation of AXB outgrowth into BC.

To deepen our understanding of the molecular mechanisms involved in AXB dormancy and activation to form a BC, we next performed the systematic analysis of transcriptomic data of the three genotypes by gene set enrichment analysis (GSEA). To this end, we used gene sets previously associated with AXB dormancy and activation in Arabidopsis and potato (Nicolas *et al*., 2022; Table S4). The genes upregulated in RdV vs *FT3^OE^* AXBs were enriched in dormancy marker genes (NES= 1.49). This was also observed, to a lesser extent in RdV vs H4 AXBs, which develop into stolon without a dormancy period (NES= 1.44). In addition, genes downregulated in RdV vs H4 were slightly enriched in activation markers (NES= -1.24) (Fig 4b; Table S5). These results point to the induction of dormancy in undifferentiated AXBs of RdV, but not of H4 and *FT3^OE^*, and to genes likely involved in the control of this process in strawberry.

As the regulatory network controlling AXB dormancy and lateral branch outgrowth converges on BRC1 (González-Grandío *et al*., 2013, 2017; Wang *et al*., 2019; Barbier *et al*., 2019), we further performed GSEA using BRC1-dependent gene sets either upregulated (BRC1up) or downregulated (BRC1down) in the dormant AXB of Arabidopsis (González-Grandío *et al*., 2013; van Es *et al*., 2024). Enrichment analysis indicated that BRC1up genes, e.g. *HB40* and *NCED3*, were also upregulated in RdV (dormant AXB) compared to H4 and *FT3^OE^* (Fig. 4b; Table S5) while BRC1down genes, e.g. ribosomal proteins S15 and S21 and fructokinase-like 1, were also downregulated in RdV compared to H4 and *FT3^OE^* (Table S5). These results, which are consistent with the known function of *BRC1* in Arabidopsis, highlight the role of *FveBRC1* in the control of strawberry AXB dormancy, through the induction of genes involved in hormone signaling and the repression of genes related to protein synthesis and metabolic activity (González-Grandío *et al*., 2017).

In addition, GSEA uncovered the previously unknown link of *FveBRC1* with AXB fate. In fact, when comparing *FT3^OE^* to H4, which present distinct fates without dormancy period, GSEA showed, no enrichment in gene set related to AXB dormancy or activation but surprisingly an enrichment in the downregulated genes of the two BRC1-dependent gene sets (BRC1up and BRC1down) in *FT3^OE^* (Fig. 4b; Table S5). Interestingly, the expression level of most BRC1down genes were higher in H4 (stolon outgrowth) than in *FT3^OE^* (BC outgrowth) (Fig. 4b; Table S5). Also unexpected is the high expression level of several BRC1up genes in H4 AXBs which will produce stolons without any dormancy stage (Fig. 4b; Table S5), suggesting that BRC1 plays a prominent and hitherto unknown role in the regulation of AXB differentiation into stolon.

### *FveBRC1* contributes to AXB fate determination and dormancy in strawberry

Given the central role of *BRC1* in repressing AXB outgrowth (Aguilar-Martinez *et al*., 2007; Wang *et al*., 2019) and its possible role in the determination of AXB fate, which has never been investigated to date, we further explored the function of *FveBRC1* in strawberry. To strengthen the association between *FveBRC1* and branching in strawberry using a different genetic background, we first exploited the previously generated ‘Ilaria’ population (Tenreira *et al*., 2017) by deriving four S5-homozygous mutants combining mutations in *FveTFL1* and *FveGA20ox4*, two genes whose role in AXB fate is well established (Iwata *et al*., 2012; Tenreira *et al*., 2017) (Fig. 5a,b). We analyzed *FveBRC1* expression in undifferentiated AXBs collected at node 4 at 4 leaf-stage as for our previous transcriptomic analysis. Results clearly showed the higher abundance of *FveBRC1* transcripts in stolon-producing AXBs (Ilaria S5-1) than in BC-producing AXBs (Fig. 5c). Nevertheless, the abundance of *FveBRC1* transcripts is likely independent of the allelic status of *FveTFL1* and *FveGA20ox4* in the ‘Ilaria’ population (Fig. 5c).

**Fig. 5.**
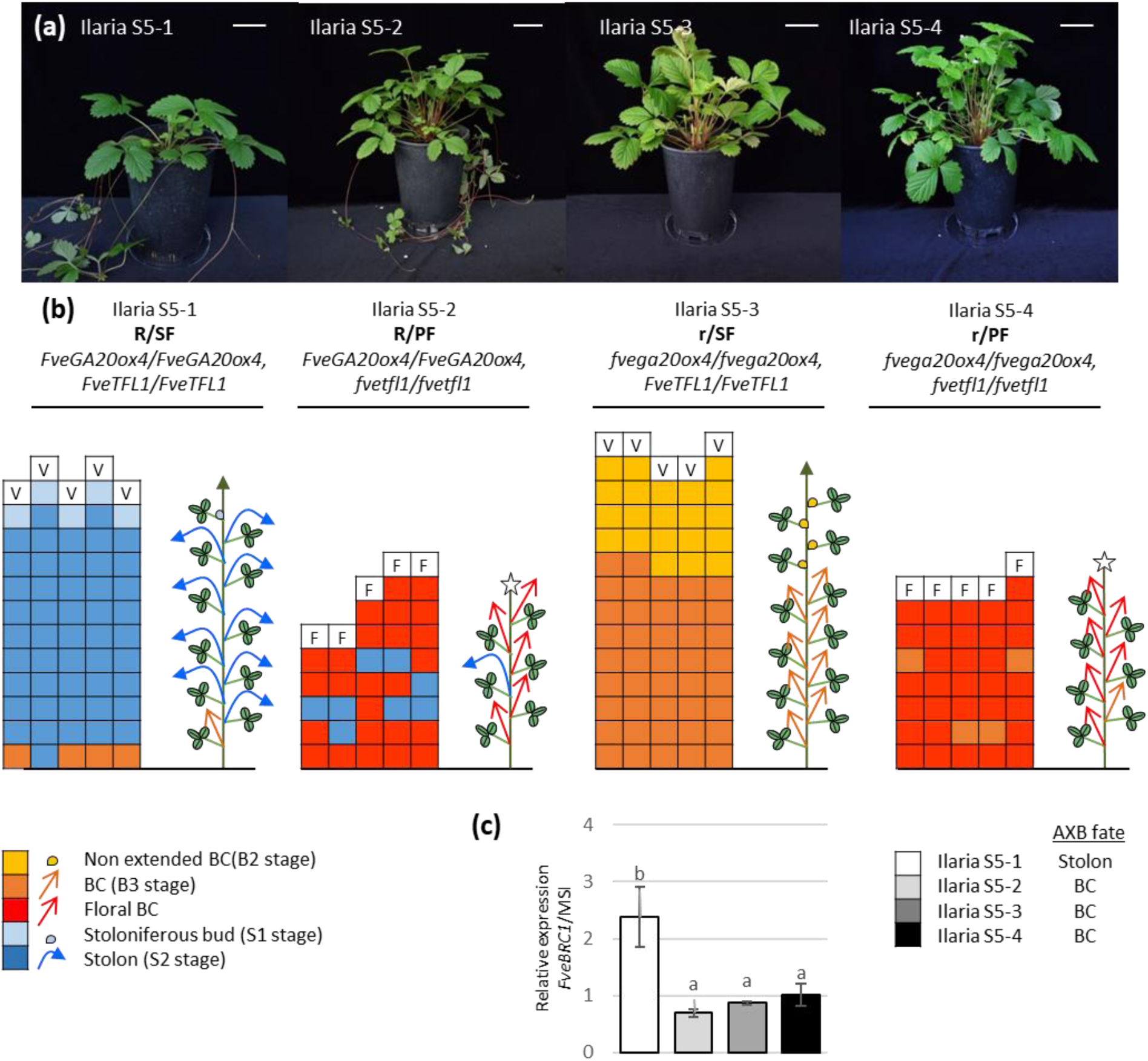
‘Ilaria’ population phenotyping and *FveBRC1* expression analysis. (a) 6-month-old plants from the ‘Ilaria’ population. White bar, 5 cm. (b) The architecture of Ilaria S5-1 to S5-4 of 6-month-old plants was determined as described in Figure 2 (n= 5). Each square box represents a node including the AXB. Box colors correspond to the stages of AXB development as described in Figure 1 legend: green, undifferentiated bud (U); pale blue, stoloniferous bud (S1); blue, stolon (S2); yellow, branch crown (BC) bud (B1); orange, dormant bud or non-extended BC (B2); dark orange, BC (B3). Red box, floral BC. V = vegetative apical meristem, F = floral apical meristem. The schematic plant architecture is shown for each genotype. (c) *FveBRC1* transcript expression of undifferentiated AXBs at node 4 of 4-leaf stage plants in the ‘Ilaria’ population. Error bars, mean + SD (n=3), with transcripts normalized to *MSI*. Significant differences were calculated by pairwise multiple comparisons using Kruskal-Wallis test followed by a Tukey post hoc test and indicated by a letter.

After confirming the association of *FveBRC1* transcript level with stolon phenotype in an independent genetic background, we then examined by *in situ* hybridization the spatial accumulation of *FveBRC1* transcripts in AXBs at the 4-leaf stage in H4, *FT3^OE^* and RdV (Fig. 6a). *FveBRC1* transcripts were much less abundant in *FT3^OE^* AXBs, where BC outgrowth occurs rapidly, than in H4 AXBs, which produce stolons, and in RdV AXBs, which produces dormant AXBs. These observations reinforce the transcriptomic findings (Fig. 4a; Table S2) that a decrease in *FveBRC1* expression is associated with AXB outgrowth into BC and determination of AXB fate. To further investigate this hypothesis, we generated mutant lines using CRISPR/Cas9 gene editing in the H4 genotype by targeting the exon 1 of *FveBRC1,* which contains the TCP domain with two guide RNAs (Fig. 6b). We selected four T1-homozygous allelic mutants. Two mutants displayed frameshift mutations, including a deletion of one base in both guide sequences in the *CR-fvebrc1#1* mutant, and an insertion of one base in guide 2 in the *CR-fvebrc1#4* mutant. *CR-fvebrc1#2* and *CR-fvebrc1#3* mutants had a large deletion and an in-frame 3 pb deletion, respectively. These mutants displayed a pronounced bushy phenotype with no stolon compared to H4 (WT) (Fig. 6c). The bushy phenotype was due to the presence of a high number of BCs and their associated trifoliate leaves, both of which increased drastically over the 18 months of observation (Fig. 6d,e). Plant architecture analysis further showed that the modification of the BC/stolon balance is due to a change in AXB fate determination as all AXBs developed into BCs, instead of stolons as in H4 (Fig. S4). Remarkably, even the AXBs in the axils of the juvenile leaves at the first two nodes developed into BCs, whereas they remained dormant in H4 (Fig. S4). Thus, the bushy phenotype of these mutants is also due the continuous development of BCs on the successive orders of branching (secondary, tertiary etc. BCs) (Fig. S4). As the mutation of *FveGA20ox4*, a key AXB-localized GA biosynthesis enzyme that is down-regulated in the BC-forming genotypes (Fig. 4a), triggers the formation of stolons at the expense of BCs (Tenreira *et al*., 2017), we further assessed its expression in AXBs of *CR-fvebrc1#2*. *FveGA20ox4* expression was largely reduced in the mutant (Fig. S5), thus suggesting that the control of stolon formation by *FveBRC1* can be mediated by *FveGA20ox4*.

**Fig. 6.**
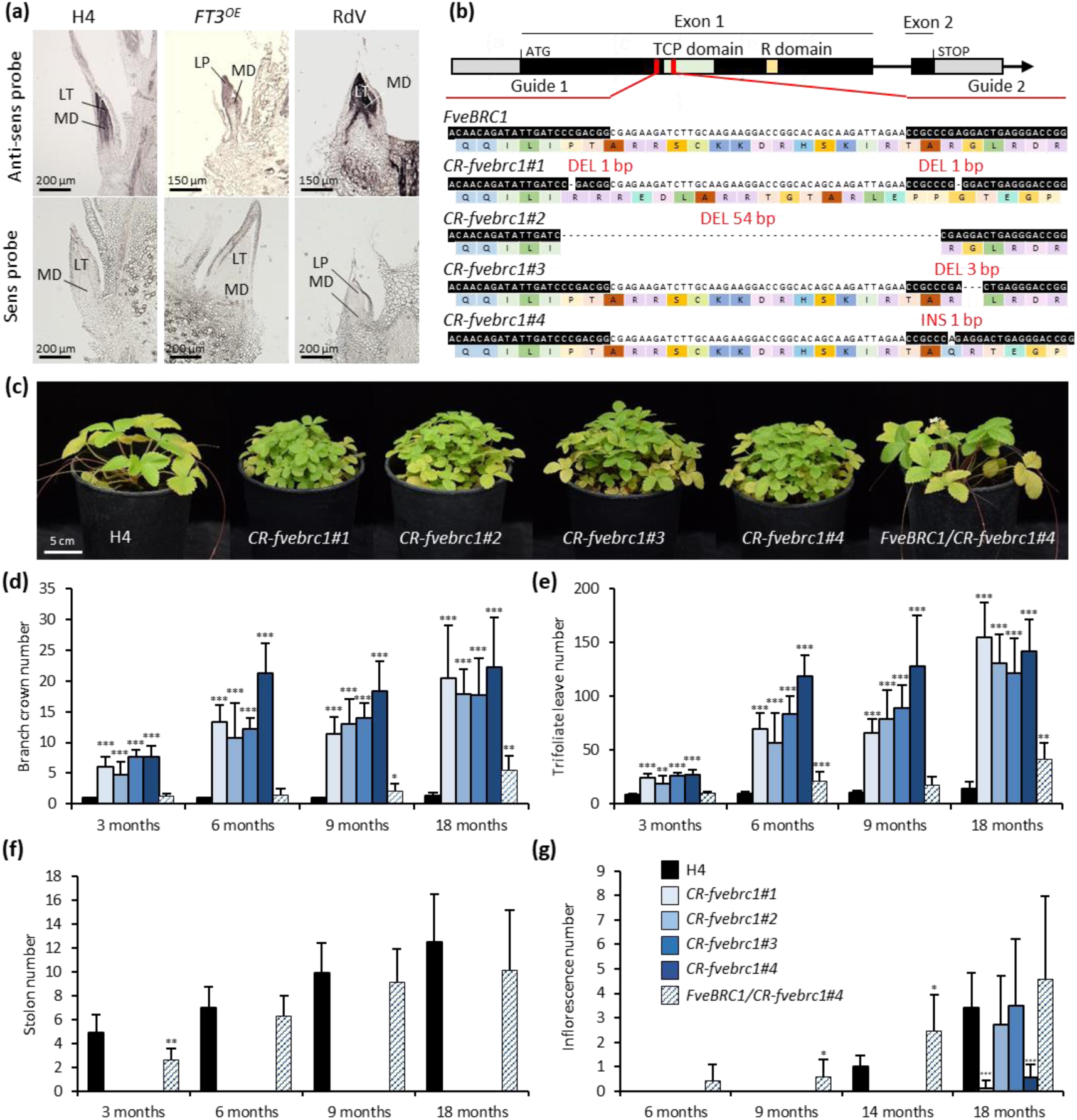
*FveBRC1* modulates plant architecture by acting on AXB fate determination and AXB outgrowth. (a) *In situ* hybridization of *FveBRC1* on undifferentiated AXBs of 4-leaf stage plants of three genotypes ‘Hawaii-4’ (H4), *35S::FveFT3^FveOE^* (*FT3^OE^*) and ‘Reine des Vallées’ (RdV). MD, meristematic dome; LT, leaflet; LP, leaf primordium. (b) CRISPR/Cas9-induced mutations in exons of *FveBRC1* produced four independent mutations. Red lines indicate the two guide RNA target sequences. Insertions (INS) or deletion (DEL) are indicated in red. (c) 10-month-old plants of four T1-homozygous *CR-fvebrc1* mutants and the T1-heterozygous mutant in comparison with the wild-type H4. (d-g) Bar graphs represent the number of (d) branch crown, (e) trifoliate leaf, (f) stolon and (g) inflorescence in H4 and *CR-fvebrc1* mutants. Wilcoxon-Mann-Whitney test showed significant differences (*, p≤0.05; **, p≤0.01; ***, p≤0.001) between H4 and the mutants (n=10).

In addition, we analyzed a T1-heterozygous mutant *FveBRC1/CR-fvebrc1#4*. The T1-heterozygous mutant produced a number of stolons comparable to the WT (Fig. 6f) and a number of BCs and trifoliate leaves intermediate between those of T1-homozygous mutants and WT (Fig. 6d,e). While flowering was significantly delayed in the T1-homozygous *CR-fvebrc1* mutants compared to the WT, the T1-heterozygous mutant flowered eight months earlier than the WT (Fig. 6g). Preliminary results further indicated that the fruit production of the heterozygous mutant could be increased by two-fold.

## Discussion

In *Fragaria* species, AXBs located in the leaf axil on the stem (primary crown) can differentiate into either lateral shoot bearing inflorescences (BC) or specialized stems (stolons), which produce daughter-plants, enabling strawberry plant multiplication. Controlling AXB fate is therefore essential, as it determines the balance between sexual and asexual reproductions, and hence the antagonistic yields of fruit and daughter plants in strawberry (Tenreira *et al*., 2017). AXB fate has been shown to be controlled by environmental signals such as day length and temperature (Andrés *et al*., 2021; Andrés and Koskela, 2022), and by endogenous signals including GA (Thompson and Guttridge, 1959; Hytönen *et al*., 2009; Tenreira *et al*., 2017; Caruana *et al*., 2018) and cytokinins (Pritts *et al*., 1986; Li *et al*., 2021; Han *et al*., 2024). However, a comprehensive view of the determination of AXB fate at molecular level is still lacking.

Potato, a well-studied stolon-forming crop species, is often used as a reference for studying AXB in strawberry (Gaston *et al*., 2020) since, depending on their position, potato AXBs have the potential to remain dormant or develop into a branch or a stolon (Nicolas *et al*., 2022). To date, major advances in potato, Arabidopsis and other plant species provided new insights into the regulation of AXB outgrowth by gene regulatory networks in response to various genetic and environmental cues (Wang *et al*., 2018; van Es *et al*., 2024). These studies have highlighted the key role of the master transcription factor *BRC1* in orchestrating AXB dormancy establishment and its subsequent release allowing AXB outgrowth into a lateral branch. However, how BRC1 could play a role in stolon differentiation has been little studied in species with dual AXB fate. In potato, technical limitations, i.e. the current inability to predict the AXB fate (stolon or lateral branch) shortly after initiation, have so far prevented transcriptomic studies from addressing the molecular mechanisms underlying the trade-off between stolon and lateral branch formation.

Here, in strawberry, by first predicting the AXB fate while still morphologically undifferentiated, and then using RNA-seq on undifferentiated AXBs from three strawberry genotypes with contrasted stolon or BC production, we were able to (i) uncover transcriptional signatures associated with different AXB fates, (ii) demonstrate that undifferentiated AXB can already recapitulate, at least partially, the developmental program leading to stolon production or BC formation and flowering and (iii) show that BRC1 not only orchestrates AXB dormancy/activation for BC production, as in other plant species studied to date, but also likely plays a hitherto unknown role in the control of stolon production. This study thus provides a comprehensive view of the molecular control of AXB fate in strawberry and highlights the dual function of BRC1 in both BC dormancy/outgrowth and stolon determination.

### The fate of undifferentiated AXBs can be predicted according to their node position on primary crown

Starting from undifferentiated AXBs, we first described their developmental trajectories according to their differentiation into either stolons or BCs (Fig. 1), as previously done for strawberry SAM (Jahn and Dana, 1970, Gaston *et al*., 2021) and, more recently, AXBs (Han *et al*., 2024). We showed that these developmental trajectories vary as a function of genotype and node position on the stem, consistent with our previous study of octoploid strawberry (Labadie *et al*., 2023b). We observed that AXBs producing stolons soon undergo basal elongation, in agreement with previous findings (Savini *et al*., 2005; Han *et al*., 2024). In contrast, AXBs producing BCs enlarge as their first leaves emerge, followed by a period of AXB dormancy (Labadie *et al*., 2023b; Han *et al*., 2024). Dormant AXBs, which develop into BCs (Han *et al*., 2024), eventually burst after a lag period of variable length. In *FT3^OE^*, a distinct H4-derived genotype producing BCs, BC formation occurs without AXB dormancy (Gaston *et al*., 2021), suggesting very early AXB activation. Thanks to this founding work on three genotypes with distinct plant architectural behaviors, producing either stolons (H4), BCs (RdV) or BCs without an intermediate AXB dormancy stage (*FT3^OE^*) (Gaston *et al*., 2021), we were able to predict the fate of a given undifferentiated AXB with high confidence.

### Undifferentiated AXBs already display transcriptional signatures related to their fate

A major finding of our study is that transcriptional signatures of undifferentiated AXBs are already very contrasted depending on their fate, despite the absence of visible morphological changes at this developmental stage. The most striking example is *FT3^OE^* where the gene expression signature of the undifferentiated AXB already includes a large number of highly expressed flowering-associated genes such as *FveFUL*, *FveLFY*, *FveFD* and *FveAP1* (Freytes *et al*., 2021), but low expression of dormancy-associated genes, including the ABA biosynthesis and signaling genes *NCED3* and *HB40* (Gonzalez-Grandio *et al*., 2017). This pattern effectively recapitulates the progression of BC differentiation and development in this genotype. Conversely, the undifferentiated AXB of the BC-forming genotype RdV, which will undergo dormancy after bud enlargement, already exhibits high expression of dormancy-related genes. These include the signal integrator *FveBRC1,* which promotes AXB dormancy, the ABA-related genes *NCED3* and *HB40,* and the strigolactone biosynthesis gene *MAX1*, which reinforces establishment of bud dormancy by enhancing *BRC1* activity (Gomez-Roldan *et al*., 2008; Kelly *et al*., 2023; van Es *et al*., 2024). Additionally, the *FveMYB117a*-regulated *CKX1* gene, which represses cytokinin accumulation and thereby prevents AXB outgrowth to BC (Han *et al*., 2024), is highly expressed in RdV AXB where it probably impedes AXB activation.

The distinction between transcriptional signatures associated with either BC or stolon fates is more complex, as the formation of both requires meristem differentiation and vascular development, which are regulated by various hormones (gibberellin, cytokinin, auxin, ethylene and brassinosteroid) and sugars (Guo *et al*., 2021; Barbier *et al*., 2019; Aloni, 2021; Yuan *et al*., 2024). Nevertheless, cluster analysis highlighted genes more specifically linked to either stolon or BC fate, most of which have unknown roles in strawberry (Fig. 3). Among the meristem-related TFs strongly up-regulated in stolon-forming AXBs is *FveBFT*, a homolog of *FveTFL1,* a systemic long-day antiflorigen contributing to photoperiodic regulation of flowering in strawberry (Li *et al*., 2019b; Gaston *et al*., 2021). Many more meristem-related TFs are markers of BC-forming AXBs, including several AP2/B3-like TFs. Among them, two members (*FvH4_1g14660* and *FvH4_1g14680*) are homologous to the Arabidopsis *VERNALIZATION1* (VRN1) gene, which represses the floral repressor *FLOWERING LOCUS C* (*FLC*) during vernalization, thus accelerating floral transition (Levy *et al*., 2002). *FveSTM-like* is homologous to Arabidopsis *STM*, a KNOTTED1-like homeobox (KNOX) gene essential for AXM initiation, acting through meristem maintenance and hormonal regulation (Hay and Tsiantis, 2010; Lechon *et al*., 2025). *FveYAB2* and *FvePIb* are homologous to Arabidopsis *YABBY* and *PISTILLATA* genes, which regulate abaxial cell fate and laminar outgrowth (Siegfried *et al*., 1999), and petal and stamen specification (Goto and Meyerowitz, 1994), respectively. Their functions are consistent with their possible role in shoot and inflorescence development in BC. We also identified the MYB TF *FveMYB117b* (*FvH4_5g27870*) as a marker of BC fate, with similar expression in *FT3^OE^* and RdV AXBs. *FveMYB117b* is a close homologous of *FveMYB117a* (*FvH4_4g31190*), recently identified as a key repressor of AXB outgrowth for BC formation (Han *et al*., 2024). However, CRISPR-editing of *FveMYB117b* had no discernible effect on AXB outgrowth, leaving its precise role in AXB fate determination unresolved.

Genes involved in hormonal biosynthesis and signaling are well represented in the cluster analysis associated with AXB fate, as expected given their role in the regulation of shoot branching (Barbier *et al*., 2019; Yuan *et al*., 2023). To date, in strawberry AXB, GA has been primarily linked to stolon formation (Tenreira *et al*., 2017; Caruana *et al*., 2018; Andrés *et al*., 2021), auxin and ABA to AXB dormancy (Qiu *et al*., 2019) like in Arabidopsis and potato (Gonzalez-Grandio *et al*., 2017; Nicolas *et al*., 2022), and CK to BC outgrowth (Qiu *et al*., 2019; Han *et al*., 2024). Among the AXB fate-related clusters, several genes encode GA 20-oxidases and GA 2-oxidases. Bioactive GAs that promote organ differentiation and growth are synthesized in meristem primordia through the key GA biosynthetic enzyme GA 20-oxidase (Tenreira *et al*., 2017). To maintain meristem indeterminacy, GA 2-oxidase converts bioactive GAs into inactive forms, preventing their activity in the meristematic dome (Galinha *et al*., 2009; Veit, 2009, Andrés *et al*., 2014; Tenreira *et al*., 2017). As expected, *FveGA20ox4*, which, when inactivated, leads to runnerless phenotype (Tenreira *et al*., 2017), was highly expressed in stolon-forming AXBs, confirming the prominent role of this specific GA20 oxidase in stolon formation. The function of GA in stolon and BC outgrowth appears more complex, as several GA 20-oxidases and GA 2-oxidases are differentially expressed in AXBs with different fates, suggesting that their specific actions depend on the precise timing and localization of their expression. Unexpectedly, BC-forming AXBs exhibited higher expression of a GA receptor, *GID1B-like,* known to target DELLA for degradation (Hauvermale *et al*., 2012) whereas the absence of the DELLA protein, FveRGA1, leads to stolon formation in a *runnerless* strawberry genotype (Caruana *et al*., 2018).

Like GAs, CKs are key regulators of plant growth and development, particularly in cell divisions. In AXB, CKs primarily regulate BC outgrowth (Waldie and Leyser, 2018; Yuan *et al*., 2023; Han *et al*., 2024). CK levels are fine-tuned by CKX (cytokinin oxidase) genes, which catalyze CK degradation (Schmülling *et al*., 2003) and are differentially expressed in stolon- *vs.* BC-forming AXBs (Fig. 4; Table S2). The high expression of a *LOG* gene, *FveLOG3*, in AXBs of BC-forming genotypes suggests that cytokinin activation is required not only for BC outgrowth (Han *et al*., 2024) but also at an earlier stage, *e.g.* at the bud enlargement (B1) stage.

These findings highlight the early specialization of undifferentiated AXBs. Although no morphological changes are visible at this stage, a specific developmental trajectory is already programmed through a distinct network of transcription factors and hormones cross-talks. This regulatory network likely plays a crucial role in AXB fate determination, with GA playing a prominent role in AXB differentiation into a stolon and cytokinin influencing AXB differentiation into a BC (Fig. 7).

**Fig. 7.**
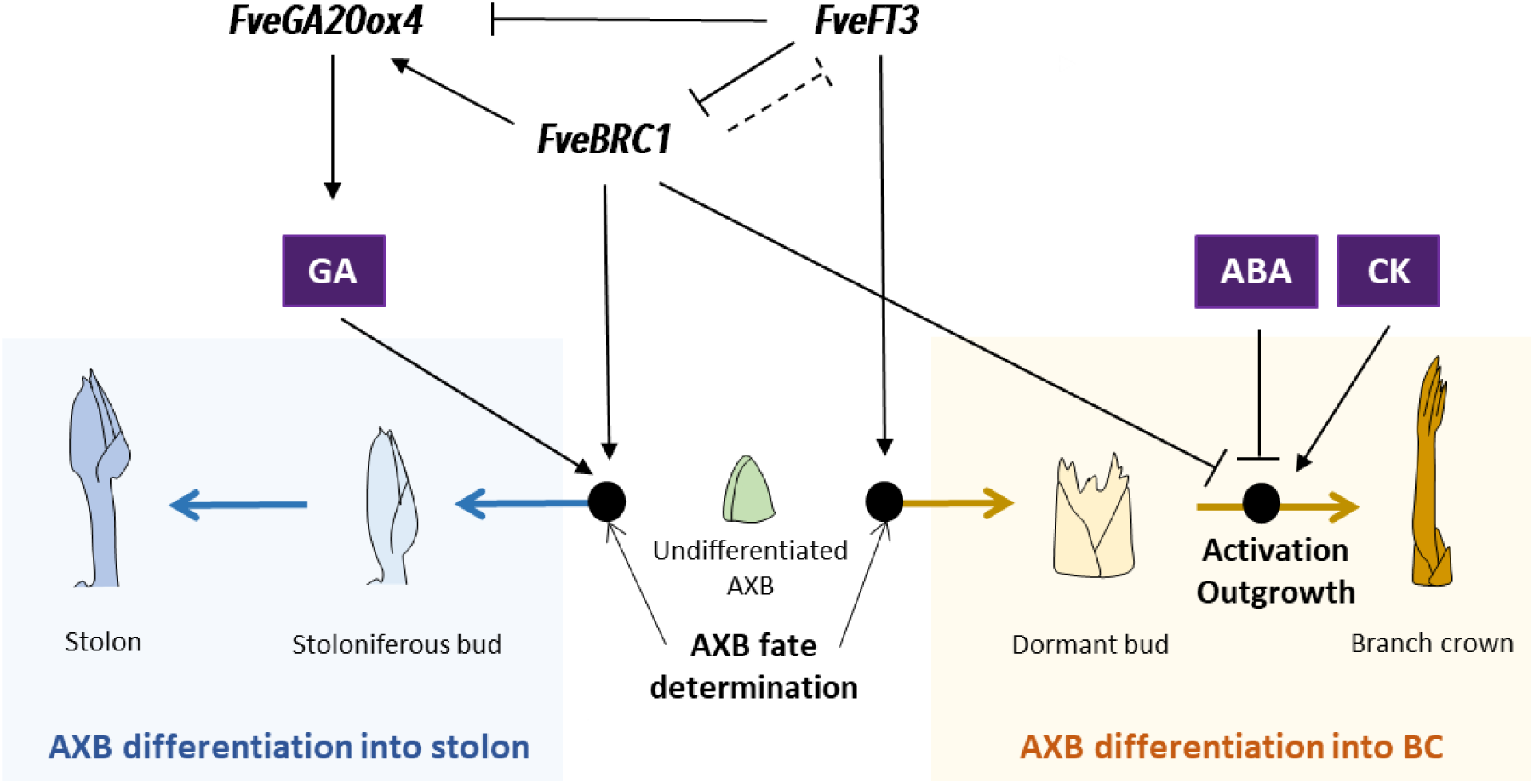
Proposed regulatory model for the determination of axillary bud fate and outgrowth into stolon and branch crown in strawberry. *FveBRC1* plays a key role in axillary bud (AXB) determination into stolon. *FveBRC1* affects the expression of the gibberellin (GA) biosynthesis gene *FveGA20ox4* which promotes stoloniferous bud formation. *FveBRC1* may also play a role in AXB fate determination via its interaction with *FveFT3*, which promotes the production of branch crowns (BCs) by repressing the expression of *FveGA20ox4*. *FveBRC1* also acts as a repressor of AXB outgrowth into BC. Activation of the dormant bud into BC is also inhibited by abscisic acid (ABA) while cytokinin (CK) promotes BC outgrowth. Arrows indicate activation and bars repression. Dotted line is hypothetical.

### *FveBRC1* controls not only AXB outgrowth into branch crowns but also AXB fate in strawberry

Our study, based on both transcriptomic analysis of undifferentiated AXBs and functional validation *in planta*, confirms that *FveBRC1* induces AXB dormancy and represses shoot branching in strawberry, supporting previous indications on its role in BC outgrowth in the strawberry *FveMYB117a* mutant (Han *et al*., 2024). The role of BRC1 as a key regulator of shoot architecture by repressing AXB outgrowth has already been demonstrated in many plant species (Aguilar-Martínez *et al*., 2007; Poza-Carrión *et al*., 2007; Braun *et al*., 2012; Muhr *et al*., 2018; Wang *et al*., 2019). An original finding of our study is that *FveBRC1* likely fulfils an additional role in strawberry by being involved in the control of AXB fate determination. Such a role has been proposed in potato, where the two paralogs *StBRC1a* and *StBRC1b* control stolon/tuberization and branch formation (Nicolas *et al*., 2015, 2022). Remarkably, the expression of *FveBRC1* is even higher in H4 (stolons) than in RdV (dormant AXBs/BCs) suggesting that the tight repression of AXB outgrowth to BC by *FveBRC1* is indeed required for stolon formation. Two other TCP genes (*FveTCP11* and *FveTCP18*) also show higher expression in H4 than in *FT3^OE^* or RdV, suggesting their involvement with *FveBRC1* in stolon fate determination. Thus, two different developmental trajectories are possible for undifferentiated AXBs displaying high *FveBRC1* expression: (*i*) AXB dormancy, eventually followed after a lag period of variable length by AXB activation and BC outgrowth, and (*ii*) stolon outgrowth.

How can *FveBRC1* specifically regulate stolon outgrowth? A possible clue is that CRISPR-engineered *FveBRC1* mutations lead to the reduction of *FveGA20ox4* expression in AXB (Fig. S5), suggesting that high *FveBRC1* activity is required for maintaining high *FveGA20ox4* expression. Moreover, the expression patterns of *FveBRC1* (Fig. 6A) and *FveGA20ox4* (Tenreira *et al*., 2017) in meristematic dome of stolon-forming AXBs are similar. As the GA biosynthesis enzyme *FveGA20ox4* and the GA signaling repressor *FveRGA1* (DELLA) are key in stolon fate determination in strawberry (Tenreira *et al*., 2017; Caruana *et al*., 2018; Li *et al*., 2018), we may speculate that *FveBRC1* effect on stolon differentiation is driven through the GA biosynthesis and regulatory network. Moreover, in Arabidopsis, different GA-associated genes are directly regulated by *AtBRC1* (van Es *et al*., 2024) and, in potato, the *StAST1* TCP gene induces the expression of *StGA20ox1* during tuberization (Sun *et al*., 2024), leading to the hypothesis that *FveGA20ox4* is a potential target of *FveBRC1*. Conversely, the effect of changes in the ratio of GA to other hormones in stolon-producing AXBs (Fig. 5) on the modulation of *FveBRC1* expression may also be considered (Liang *et al*., 2022).

*FveBRC1* could also play a role in AXB fate determination via its interaction with *FveFT3*, which promotes the production of BCs instead of stolons (Gaston *et al*., 2021), a role opposite to that of *FveBRC1.* We further showed here that *FveFT3* overexpression leads to the reduction of *FveBRC1* expression. In addition, protein-protein interactions between BRC1 and FT family members have been shown in various species (Niwa *et al*., 2013; Maurya *et al*., 2020; Feng *et al*., 2022, Nicolas *et al*., 2022), suggesting competition between the two. In addition, in potato, StBRC1B induces AXB dormancy by interacting with the tuberigen protein SP6A, from the FT/TFL1 family, to block tuber induction activity in the aerial nodes (Navarro *et al*., 2011; Nicolas *et al*., 2022).

In conclusion, our findings indicate that AXB fate is established very early after AXB initiation and provide gene targets for studying the developmental processes underlying stolon and BC formation as well as their regulation by various endogenous cues, including the BRC1 repressor. Our study and published results suggest that BRC1 not only plays a central function as a negative regulator of AXB branch outgrowth, by keeping AXB dormant, but also plays at the same time an essential but less well-established role in enabling AXB stolon outgrowth (Fig. 7). Future developments will take advantage of our large gene expression dataset to explore the mechanisms involved in AXB differentiation, in particular the novel and unique function of *FveBRC1* in stolon fate determination. From a more applied perspective, it will be worth exploring the modulation of *FveBRC1* activity by gene editing for improving strawberry production period and yield, as suggested by the preliminary results obtained with the heterozygous mutant.

## Acknowledgments

The research work was carried out thanks to the support of INRAE BAP MeriFate project and Nouvelle-Aquitaine AgirClim project (Grant numbers 2018-1R20202).

## Author Contributions

MA, BD and AG conceived the studied and designed experiments. MA organized experiments and MA, PP, AP and MH realized experiments. MA realized the histological, statistical and bioinformatic analyses with the help of YC and PM. AG, BD, CR and MA wrote the manuscript with the help of MN. All the authors discussed the results and commented the manuscript.

## Data availability

All data can be found within the manuscript and its supporting information.

## Supporting Information

**Fig. S1. ‘Hawaii-4’ (H4), *35S::FveFT3^FveOE^* (*FT3^OE^*) and ‘Reine des Vallées’ (RdV) plants at the 5-leaf stage.**

**Fig. S2. Morphological characterization of AXB development into a branch crown in ‘Reine des Vallées’ (RdV).**

**Fig. S3. Phylogenetic analysis of TCP proteins.**

**Fig. S4. Schematic representation of the architecture of 4-month-old ‘Hawaii-4’ (H4) and *CR-fvebrc1* T1 plants.**

**Fig. S5. Expression pattern of *FveGA20ox4* in *CR-fvebrc1#2* undifferentiated AXBs.**

**Table S1. List of primers used in the manuscript.**

**Table S2. Table S2. Results of RNAseq analyses.**

**Table S3. Significant Gene Ontology (GO) categories (p-value < 0.05) for each cluster.**

**Table S4. Gene sets used for gene set enrichment analysis (GSEA).**

**Table S5. Gene Set Enrichment Analysis (GSEA) results for each pairwise comparison.**

